# A Clear, Legible, Explainable, Transparent, and Elucidative (CLETE) Binary Classification Platform for Tabular Data

**DOI:** 10.1101/2023.06.20.545752

**Authors:** Ahmad Nasimian, Saleena Younus, Özge Tatli, Emma U. Hammarlund, Kenneth J. Pienta, Lars Rönnstrand, Julhash U. Kazi

## Abstract

Therapeutic resistance continues to impede overall survival rates for those affected by cancer. Although driver genes are associated with diverse cancer types, a scarcity of instrumental methods for predicting therapy response or resistance persists. Therefore, the impetus for designing predictive tools for therapeutic response is crucial and tools based on machine learning open new opportunities. Here, we present an easily accessible platform dedicated to Clear, Legible, Explainable, Transparent, and Elucidative (CLETE) yet wholly modifiable binary classification models. Our platform encompasses both unsupervised and supervised feature selection options, hyperparameter search methodologies, under-sampling and over-sampling methods, and normalization methods, along with fifteen machine learning algorithms. The platform furnishes a k-fold receiver operating curve (ROC) - area under the curve (AUC) and accuracy plots, permutation feature importance, SHapley Additive exPlanations (SHAP) plots, and Local Interpretable Model-agnostic Explanations (LIME) plots to interpret the model and individual predictions. We have deployed a unique custom metric for hyperparameter search, which considers both training and validation scores, thus ensuring a check on under or over-fitting. Moreover, we introduce an innovative scoring method, NegLog2RMSL, which incorporates both training and test scores for model evaluation that facilitates the evaluation of models via multiple parameters. In a bid to simplify the user interface, we provide a graphical interface that sidesteps programming expertise and is compatible with both Windows and Mac OS. Platform robustness has been validated using pharmacogenomic data for 23 drugs across four diseases and holds the potential for utilization with any form of tabular data.

## Introduction

Predicting drug sensitivity is an integral aspect of personalized medicine, with the potential to significantly alter therapeutic strategies, notably within oncology (Rafique, et al., 2021; Rodriguez and Pennington, 2018). Through the application of biological information and sophisticated computational algorithms, it becomes possible to ascertain the optimal therapeutic agent, thereby reducing unnecessary toxicity due to non-selective chemotherapy. Numerous molecular signatures, such as genomic, transcriptomic, metabolomic, and proteomic profiles, are capable of describing the unique pathophysiological states of individuals. Thus, each of these molecular descriptors, either singularly or in concert, can be deployed to project drug sensitivity by establishing correlations with drug response data. However, these molecular signatures embody a multitude of features, rendering it a challenging task for biologists to establish connections with drug response data.

Machine learning algorithms have been extensively deployed in the scientific domain to establish correlations between biological attributes and drug responsiveness, particularly in experimental and preclinical settings (Rafique, et al., 2021). Most of the studies focusing on drug sensitivity prediction have relied on a standard suite of features to forecast the reactivity of a comprehensive range of therapeutics across various disease conditions. Although these studies establish a broad analytical framework, they often lack the precision and power necessary for clinical applicability. The reactivity of specific pharmaceutical compounds is contingent on an exclusive array of biological phenomena, which must be incorporated into the feature set for effective prediction specific to both drug and disease conditions. Recent research endeavors utilizing disease-centric and drug-specific biological attributes have exhibited notable predictive efficacy (Nasimian, et al., 2023; Nasimian, et al., 2023; Shah, et al., 2021). Nevertheless, these methodological approaches necessitate substantial investment in terms of time and resources and demand a synergetic collaboration between researchers in the fields of biology and data science to ensure the effectiveness and rationality of the strategy.

The process of predicting drug sensitivity incorporates a multitude of computational procedures, encompassing aspects such as data preprocessing, feature extraction, data partitioning and sampling, optimization of model hyperparameters, and the employment of suitable machine learning algorithms (Rafique, et al., 2021). Moreover, comprehensive performance testing and interpretability evaluation are imperative for any algorithm after the training phase to elucidate the underpinning mechanism of its predictions. Although pharmacogenomic data generated within a laboratory environment constitute the core of the predictive model, the execution of data processing and model construction poses considerable challenges due to the intricate nature of the computational steps’ integration, requiring proficiency in a high-level programming language such as Python or R.

Aside from the inherent intricacies involved in the formulation of a machine learning model, the regulation of overfitting and underfitting continues to pose significant hurdles across various machine learning methodologies. A multitude of hyperparameter optimization algorithms predominantly target the maximal efficacy of a machine learning model, which may not manifest an equivalent predictive capacity on test instances. This phenomenon arises as a result of hyperparameter optimization algorithms directing their focus on the optimal output of the predictive model, whilst neglecting to factor in discrepancies in performance between training and validation datasets.

In the present methodological outline, we detail the design and functionality of a machine learning (ML) system tailored to suit the needs of experimental biologists lacking the necessary temporal resources to acquire proficiency in computational languages such as Python or R. We’ve engineered unique performance assessment metrics, taking into account both the efficacy of the model and the disparity between training and validation accuracy, thereby ensuring automated regulation of overfitting and underfitting instances. The core concept introduced herein is denoted as CLETE (clear, legible, explainable, transparent, and elucidative), which encapsulates the system’s primary attributes - providing a lucid, thoroughly interpretable ML model. Additionally, the platform offers graphical user interfaces for both model creation and prediction functionality.

### The structure of alphaML platform

The core of alphaML ML is composed of various modules including data management, feature selection, data normalization, data sampling, hyperparameter tuning, model training and validation, testing, and model interpretability. A similar method has been described recently (Shah, et al., 2023). The data management module initially verifies the data accessibility in a particular directory of the local storage. If absent, it retrieves example datasets from Google Drive. The retrieved data from the local storage is then utilized to select distinct features and preprocessed for model generation (Figure 1A). While not strictly essential, for pharmacogenomic datasets, such as those derived from microarray or RNAseq platforms, we advocate for the submission of normalized and log2-transformed data. The iDEP online platform encompasses numerous data normalization options, including edgeR (Ge, et al., 2018; Robinson, et al., 2010), which is instrumental in transforming raw counts. Subsequent to the preprocessing phase, a subset of data is used for hyperparameter optimization, and the optimized parameters are then forwarded to the model for subsequent evolution and performance testing using the test dataset (Figure 1B). Additionally, a distinct module has been integrated for generating predictions utilizing pre-established models (Figure 1C)

**Figure 1.**
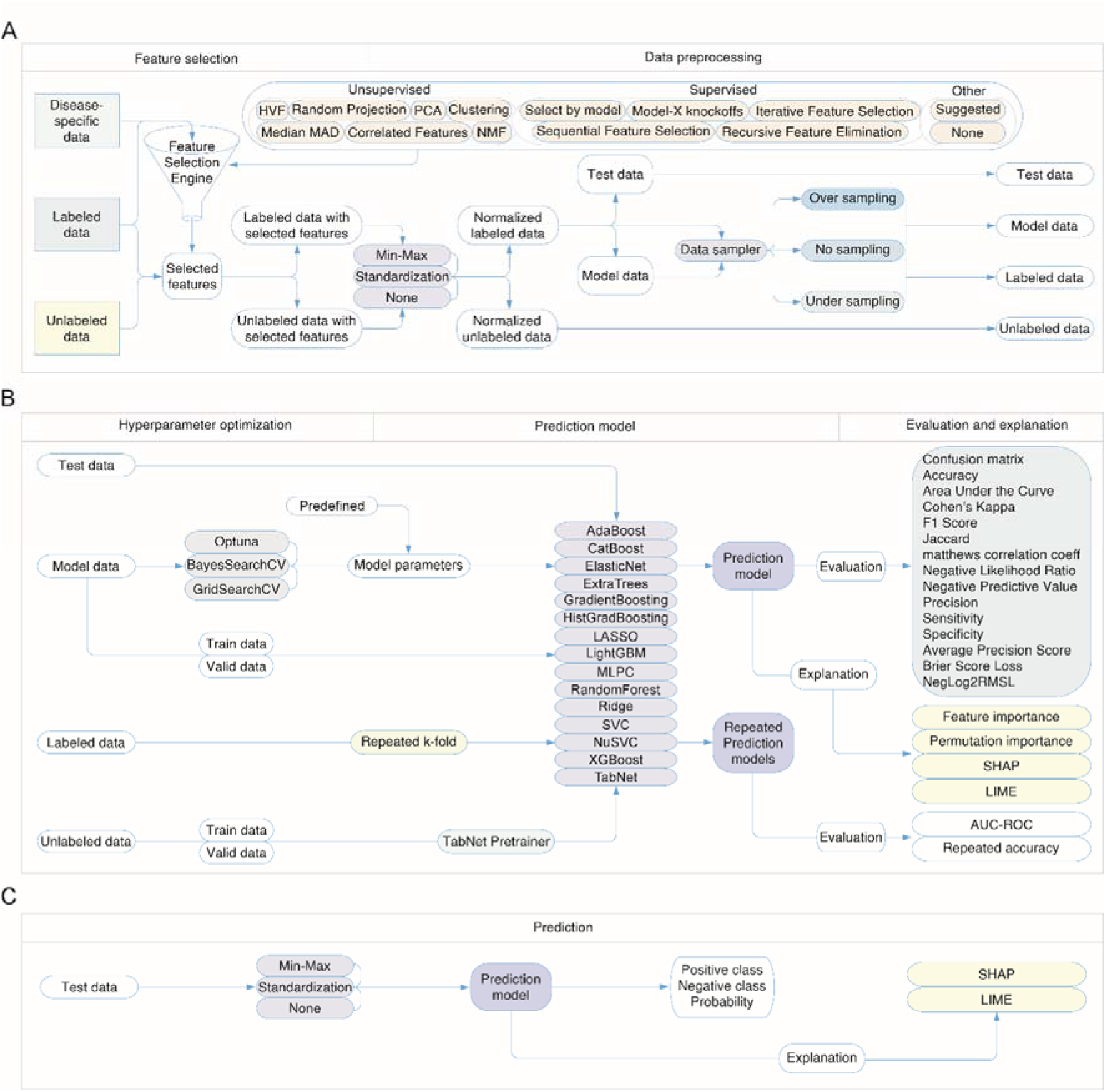
Schematic representation of the alphaML platform. A) The Data Management Process. This illustrates the process of data accessibility verification in a specific directory of local storage. It also details the process of feature selection and preprocessing of retrieved data from local storage for model generation. The flowchart signifies how the system retrieves example datasets from Google Drive in the absence of local data. B) Model Development and Evaluation. This figure visualizes the subsequent stages following data preprocessing. It showcases the use of a data subset for hyperparameter optimization and the forwarding of optimized parameters to the model for its further evolution. It also exhibits how the system tests the model’s performance using the test dataset. C) Prediction Generation. This figure depicts the functionality of an integrated module dedicated to generating predictions using pre-established models. This highlights the platform’s ability to utilize already developed models for prediction tasks, emphasizing its flexibility and adaptability.

### The feature selection module

The alphaML platform is designed to accept predefined feature lists or to implement unsupervised feature selection strategies such as the identification of Highly Variable Features (HVF), Median Absolute Deviation (MAD), Principal Component Analysis (PCA), Random Projection, Clustering, and Non-negative Matrix Factorization (NMF). Moreover, it employs supervised feature selection methodologies, encompassing model-based selection, Recursive Feature Elimination (RFE), Sequential Feature Elimination (SFE), and Model-X Knockoffs. The system is further equipped with utilities for eliminating features demonstrating correlation exceeding 90% and iterative feature selection operations - though computationally intensive - for pinpointing the top 1-3 features. The module integrates four consecutive feature selection alternatives, enabling the alphaML to conduct feature selection through diverse techniques. To exemplify, features with high variability can be identified via HVF or Median of MAD, facilitating an initial reduction in feature count prior to the execution of other techniques. Subsequent application of Random Projection, PCA, NMF, or model-based selection can further condense the feature set prior to initiating resource-demanding supervised methodologies. In circumstances where feature selection is the sole requirement, the system’s feature selection module can function independently by opting for the algorithm ‘None’.

To assess the functionality of the feature selection module, we utilized four distinct pathological conditions as exemplars: Acute Myeloid Leukemia (AML), B-cell Acute Lymphoblastic Leukemia (B-ALL), T-cell Acute Lymphoblastic Leukemia (T-ALL), and ovarian cancer. We consecutively deployed four feature selection strategies in two unique sequences: HVF -> Model-based Selection -> PCA based Selection -> Iterative Feature Selection (Figure 2A), and HVF -> Random Projection -> NMF based Selection -> Iterative Feature Selection (Figure 2B) for 23 distinct pharmaceutical compounds. Our findings revealed that features displayed a significant degree of specificity related to the pathological condition, in addition to exhibiting a level of specificity for the pharmaceutical inhibitor and feature selection methodology employed. These observations underscore the necessity for a disease and drug-specific predictive model rather than a generalized model applicable to a broad spectrum of diseases and drugs. Subsequently, we discerned an amplified synergistic outcome when Brain Expressed X-linked protein 1 (BEX1) was subject to overexpression in ovarian carcinoma cellular cultures in conjunction with erlotinib administration. This reinforces the concept that methodological selection for phenotypic features possesses significant utility in the elucidation of molecular interplay, thereby facilitating the derivation of efficacious therapeutic combinations.

**Figure 2:**
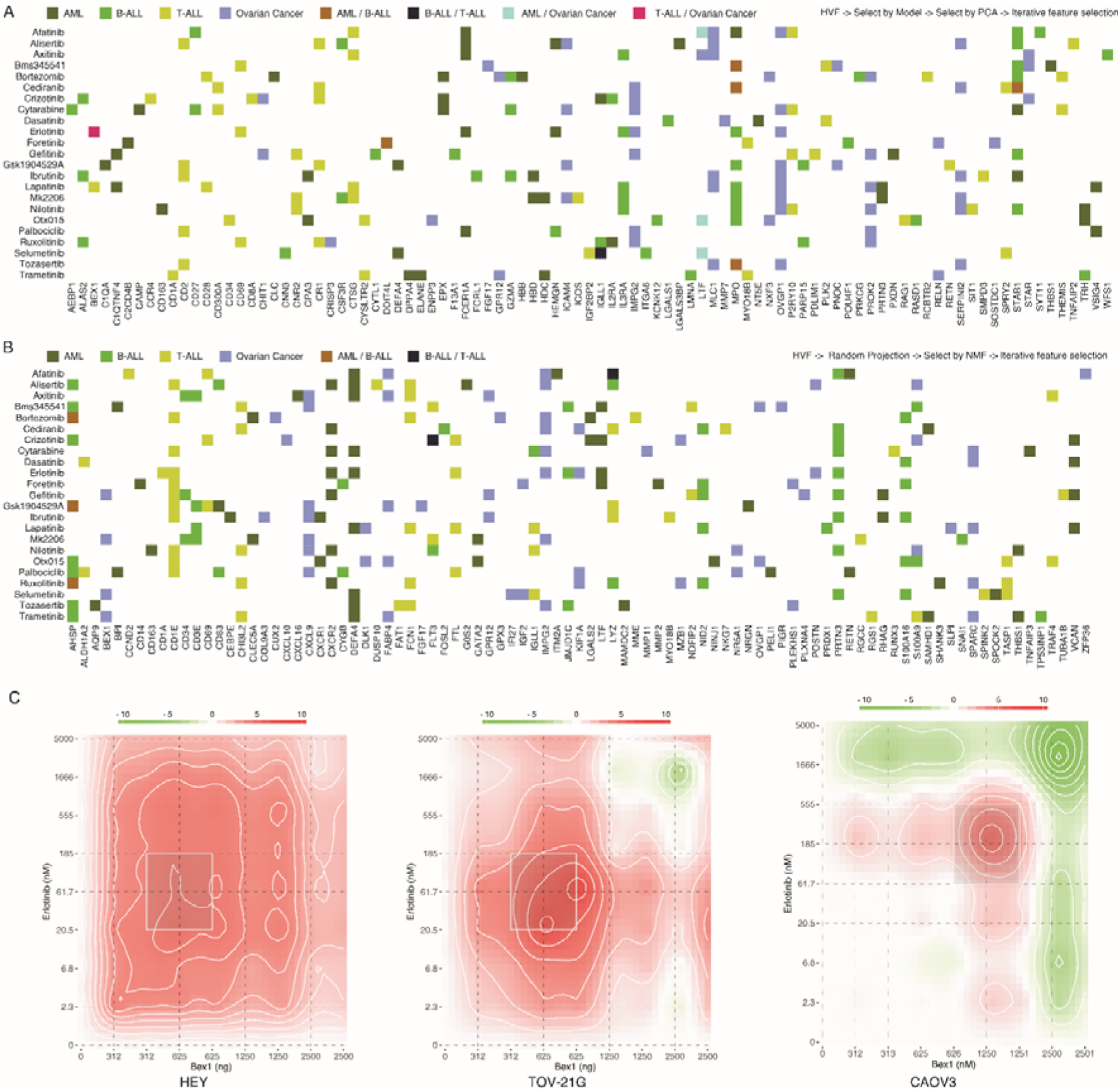
Sequential application of feature selection methods. (A) The sequential implementation of the High Variable Features (HVF), Model-based Selection (Random Forest), Principal Component Analysis (PCA) based Selection, and Iterative Feature Selection applied to AML, B-ALL, T-ALL, and ovarian cancer. (B) The sequential application of HVF, Random Projection, Non-negative Matrix Factorization (NMF) based Selection, and Iterative Feature Selection for the same diseases. A panel of 23 pharmaceutical compounds was assessed. The heatmap provides an overview of the feature selection procedures employed to assess the functionality of the feature selection module. (C) The synergistic effects of BEX1 overexpression and erlotinib treatment were quantitatively evaluated in ovarian cancer cell lines following a 48-hour treatment period. DECREASE (Ianevski, et al., 2019) was used to expand combination response and SynergyFinder (Ianevski, et al., 2020) was used to calculate the synergy.

### Data normalization module

In the realm of machine learning, the normalization of data is a critical preparatory stage that ensures equable contribution from disparate features towards the efficacy of the model, thereby augmenting the stability and efficiency of learning algorithms. In the absence of normalization, attributes exhibiting larger numerical spans may overshadow those with smaller numerical spans, potentially inducing biased model predictions. Techniques such as Min-Max scaling and Standardization are instrumental in alleviating these issues. Min-Max scaling, colloquially termed normalization, applies a transformation that coerces the data to conform to a specified range, customarily [0,1]. This is accomplished by deducting the minimum value of an attribute followed by division by the range of that attribute. This approach is advantageous when the dataset’s approximate boundaries are known, and the maintenance of precise distances between data points is nonessential. Contrastingly, Standardization, also recognized as Z-score normalization, recenters the data about the mean, culminating in a zero mean, and scales it according to the standard deviation. Consequently, the ensuing distribution possesses a standard deviation of 1. This method is relatively impervious to outliers and is typically employed when the algorithm necessitates the supposition of normally distributed data, as seen in methods involving linear regression, logistic regression, and support vector machines. Min-Max scaling and Standardization methodologies both possess their merits and applications, and the preference for one over the other relies on the specific attributes of the data and the requisites of the machine learning algorithm in use. Through the transformation of the feature space to render it more conducive to the algorithms’ presuppositions, data normalization serves as a critical determinant in the successful deployment of machine learning models. Moreover, beyond Min-Max and Standardization, alphaML platform allows an option ‘None’ to forgo any form of transformation.

### Data Sampling module

Within the domain of machine learning, specifically in relation to classification issues, techniques such as undersampling and oversampling are applied to rectify the issue of class imbalance. Class imbalance is a phenomenon where certain classes in the dataset are not as well represented as others. This disparity can create a biased model training process, with the model demonstrating superior ability in predicting the majority class and displaying subpar performance when dealing with the minority class. Undersampling mitigates this problem by diminishing the size of the majority class, thereby promoting a more equitable distribution among classes. This method entails the logical or random removal of instances from the majority class with the aim of aligning the sample size with that of the minority class. Though undersampling can effectively balance class distribution, it poses the risk of excluding potentially valuable data that could contribute significantly to model training. In contrast, oversampling attempts to rectify class imbalance by amplifying the number of occurrences within the minority class. This is typically accomplished by either replicating instances from the minority class or synthesizing new instances grounded in the extant data, the latter approach often employing a technique known as Synthetic Minority Over-sampling Technique (SMOTE). Despite its efficacy in balancing class distribution, oversampling may cause model overfitting to the minority class due to the utilization of duplicated or synthesized instances.

The imbalanced-learn library was deployed in alphaML to regulate the application of undersampling and oversampling (Lemaître, et al., 2017). Furthermore, the sampling module provides a ‘no sampling’ method that maintains the original proportions of the major and minor classes. However, during the model construction phase, a class imbalance parameter is invoked to balance the ratio. Notably, test samples always remain unaltered by the sampling process. We ensure this by partitioning the test sample prior to its submission to the sampling module, thereby guaranteeing that no sampling biases afflict the test data. Through the examination of pharmacogenomic data from 23 inhibitors and four disease models, it was observed that the ‘no sampling’ method consistently surpassed the performance of both oversampling and undersampling methods (Figure 3), intimating that the adoption of a sampling procedure may not invariably contribute towards enhancing the efficacy of a machine learning model. Nonetheless, this observation could exhibit high specificity to a particular problem, thus underscoring the perpetual necessity of a comprehensive comparative analysis.

**Figure 3.**
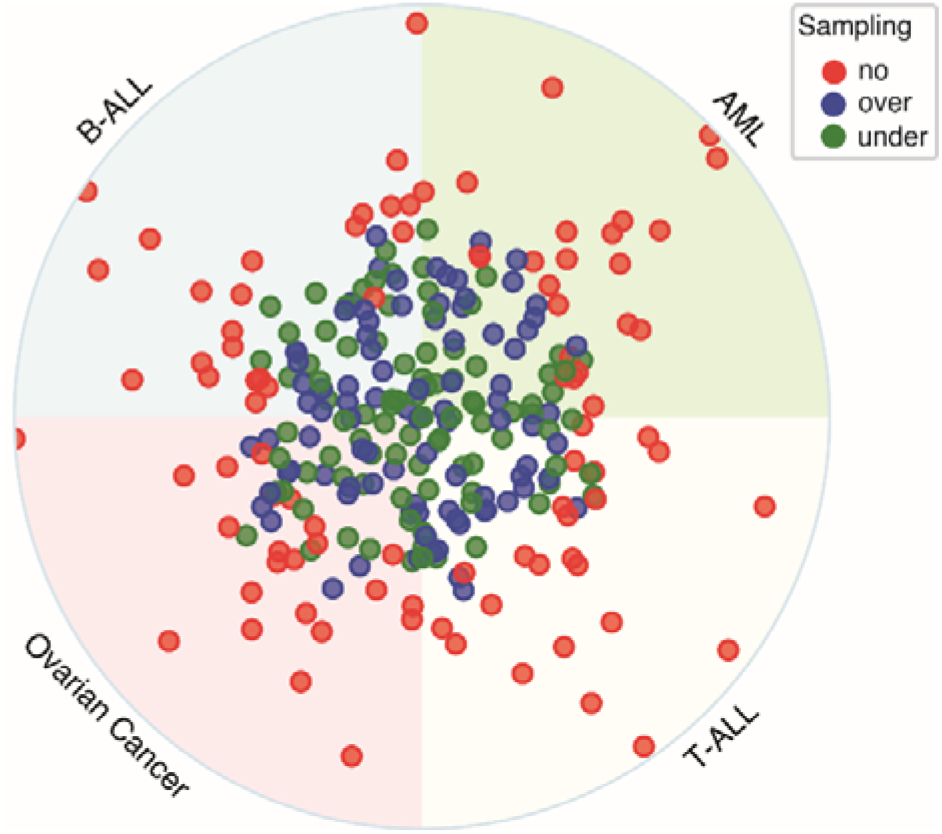
A comparative analysis of varied sampling techniques. We employed an XGBoost classifier, fixed with predetermined parameters, to construct and evaluate binary classification models. Prior to implementing undersampling or oversampling procedures, the test samples were distinctly partitioned. The radial distance of each point from the circle’s origin corresponds to the normalized NegLog2RMSL (described in Ref. (Shah, et al., 2023)) for each inhibitor, where a higher value indicates superior performance.

### Hyperparameter optimization module

The optimization of hyperparameters constitutes a fundamental stage in the development of robust and high-precision machine learning models. Hyperparameters denote factors that are not inherently derived within predictors, but rather are determined by the user preceding the model training process. Their influence on model performance is considerable as they preside over the intricacies of the model, the extent of regularization, the learning rate, among other factors. We have assimilated three varied hyperparameter tuning methodologies into our process: Grid Search Cross Validation (GridSearchCV), Bayesian Optimization, and Optuna. GridSearchCV is a conventional technique employed in hyperparameter tuning. It conducts a comprehensive exploration over a manually prescribed subset within the hyperparameter space of a learning algorithm. For each hyperparameter combination, it trains a model and gauges its performance through cross-validation, ultimately selecting the hyperparameters that offer the highest performance metric. Although it provides a comprehensive and straightforward approach, GridSearchCV can be computationally burdensome, especially when dealing with numerous hyperparameters and extensive datasets. Bayesian Optimization, implemented in libraries such as Scikit-Optimize, offers a more proficient method for hyperparameter tuning (I., et al., 2020). It constructs a probabilistic model of the objective function, leveraging previous evaluation outcomes, and employs this model to choose the hyperparameters that exhibit the highest potential for evaluation. Hence, Bayesian optimization is particularly apt for optimization problems where function evaluations impose a significant computational cost. Lastly, Optuna presents another proficient framework for hyperparameter optimization, effectively automating the hyperparameter tuning process (T., et al., 2019). It integrates several cutting-edge methods for efficacious hyperparameter exploration, such as handling of categorical parameters, pruning of unpromising trials, and automated concurrency management.

The methodology employed for hyperparameter tuning incorporated a compound scoring system that utilized both Cohen’s Kappa coefficient and the Matthew’s Correlation Coefficient (MCC). This amalgamated scoring strategy assures a rigorous evaluation of categorization models by harnessing the unique capabilities inherent in each metric.

Cohen’s Kappa, with its numerical values spanning from -1 to 1, quantifies the concurrence between two independent classifiers, effectively controlling for random chance and imbalance in class distribution. Conversely, the MCC, possessing an identical range, appraises binary categorization systems by contemplating all constituents of the confusion matrix, demonstrating resilience towards class disequilibrium. This dualistic metric strategy, exhibiting reduced sensitivity to class distribution skewness and offering an exhaustive appraisal of accuracy, fosters dependable evaluation of classifier efficacy, thus supporting superior model selection for pragmatic implementation. We employed the geometric mean of Kappa and MCC errors, notated as “kappa_mcc_error”, which is computed using the formula 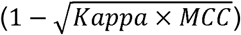. Alongside this, the “kappa_mcc_error” was utilized in conjunction with a controlled fitting approach, implemented to mitigate both overfitting and underfitting tendencies in our model. We also introduced a proprietary metric, termed “custom_score”, which incorporates the “kappa_mcc_error” in a bipartite manner, taking into account both training and validation scores.

In assessing the efficacy of hyperparameter search, we executed the Optuna hyperparameter optimization protocol. Three performance evaluation indices were exploited: the Hamming loss that signifies the converse of accuracy, “kappa_mcc_error”, and a uniquely devised metric, “custom_score”. The Hamming loss and the “kappa_mcc_error” metrics utilize validation data to generate a numerical measure, consequently guiding the model towards its optimal performance aligned with the validation score. Contrarily, the “custom_score” considers both the training and validation datasets, aiming to attenuate the divergence between the performance on training and validation while simultaneously endorsing the model’s paramount performance. In accordance with data derived from 23 inhibitors and four disease models, the incorporation of “custom_score” within the Optuna scoring architecture seems to assure superior performance as measured by NegLog2RMSL (Figure 4A). Additionally, while juxtaposing different algorithms with various search methods, “custom_score” amalgamated with Optuna exhibited the highest NegLog2RMSL (Figure 4B). In aggregate, the evidence indicates that the employment of “custom_score” can augment the performance of binary classification models.

**Figure 4.**
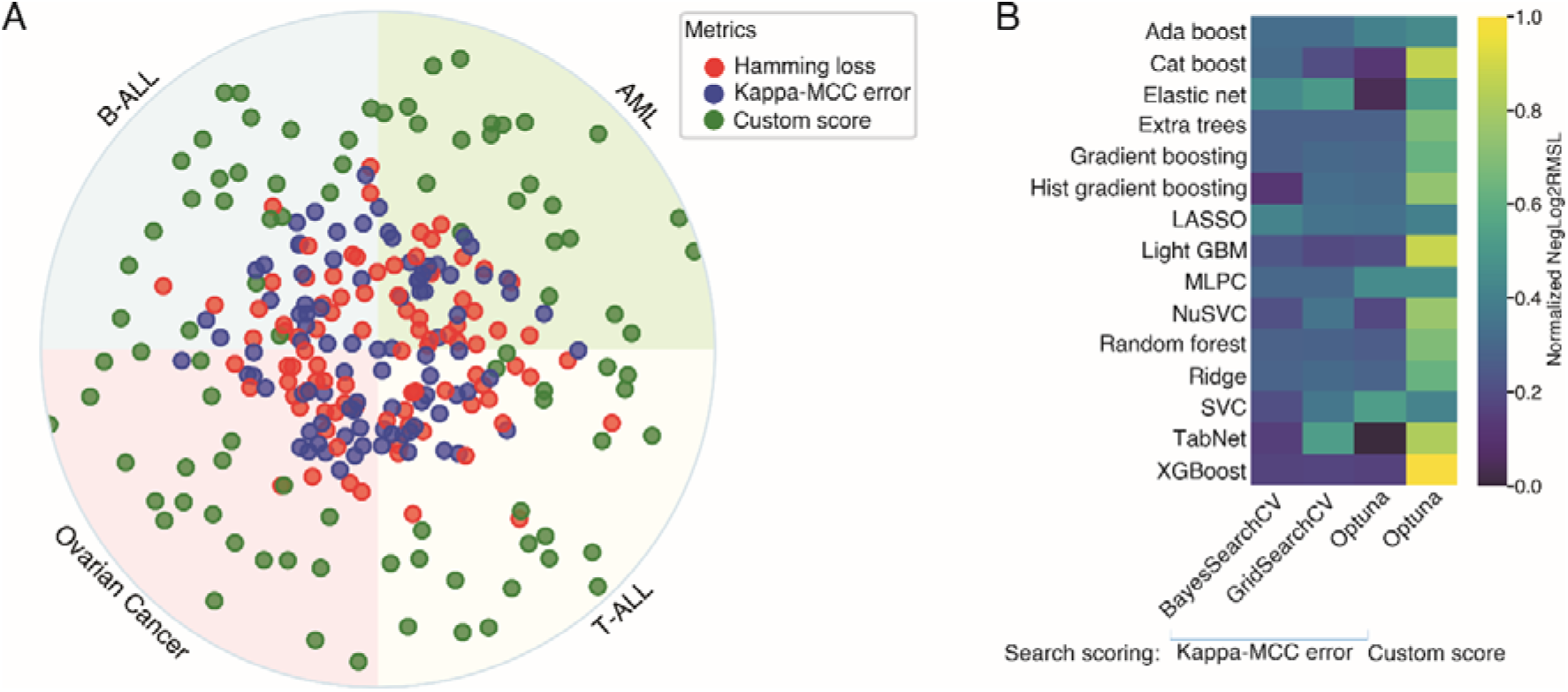
The “custom_score” exhibits superior performance in comparison to alternative scoring metrics. (A) The hyperparameters for the XGBoost classifier were optimized utilizing Optuna, encompassing three distinct scoring methods. The model incorporated four distinct disease paradigms along with 23 inhibitors. Each individual dot denotes the normalized NegLog2RMSL value, plotted in reference to the origin of the circle considered as zero. (B) Diverse search methods, which integrated either “kappa_mcc_error” or “custom_score”, were employed for the optimization of hyperparameters across a selection of algorithms. These methods were tested using T-ALL samples in conjunction with the trametinib inhibitor.

### Algorithms

The alphaML platform integrates 15 algorithms, which can be classified into four categories: regularized linear models, support vector machines, ensemble methods, and neural networks, as represented in Figure 5A. Regularized linear models serve as modifications of the standard linear regression model, integrating a regularization parameter to inhibit overfitting and augment the predictive capacity of the model across diverse data sets. These models are typically formulated to include a penalty term within the loss function, which minimizes both the residuals and the complexity of the model (Tibshirani, 1996). There are three main types of regularized linear models: ridge, lasso, and elastic net. The ridge model introduces an L2 penalty, which is equal to the square of the coefficient magnitudes scaled by a constant alpha (Hoerl and Kennard, 1970). This results in the attenuation of coefficient values; however, it does not drive them to zero, thereby not performing feature selection. The least absolute shrinkage and selection operator (lasso) integrates an L1 penalty, which is equivalent to the absolute magnitude of the coefficients (Tibshirani, 1996). This type of regularization may induce zero coefficients, thereby performing both parameter shrinkage and feature selection. The Elastic Net model serves as a hybrid methodology, integrating both L1 and L2 penalties (Zou and Hastie, 2005). It is notably advantageous when multiple features are correlated. By merging aspects from both ridge and lasso, it retains the feature selection capabilities of lasso, while maintaining the regularization properties of ridge. These regularization techniques have been substantiated to be successful across a diverse range of applications, demonstrating both predictive accuracy and model interpretability.

**Figure 5.**
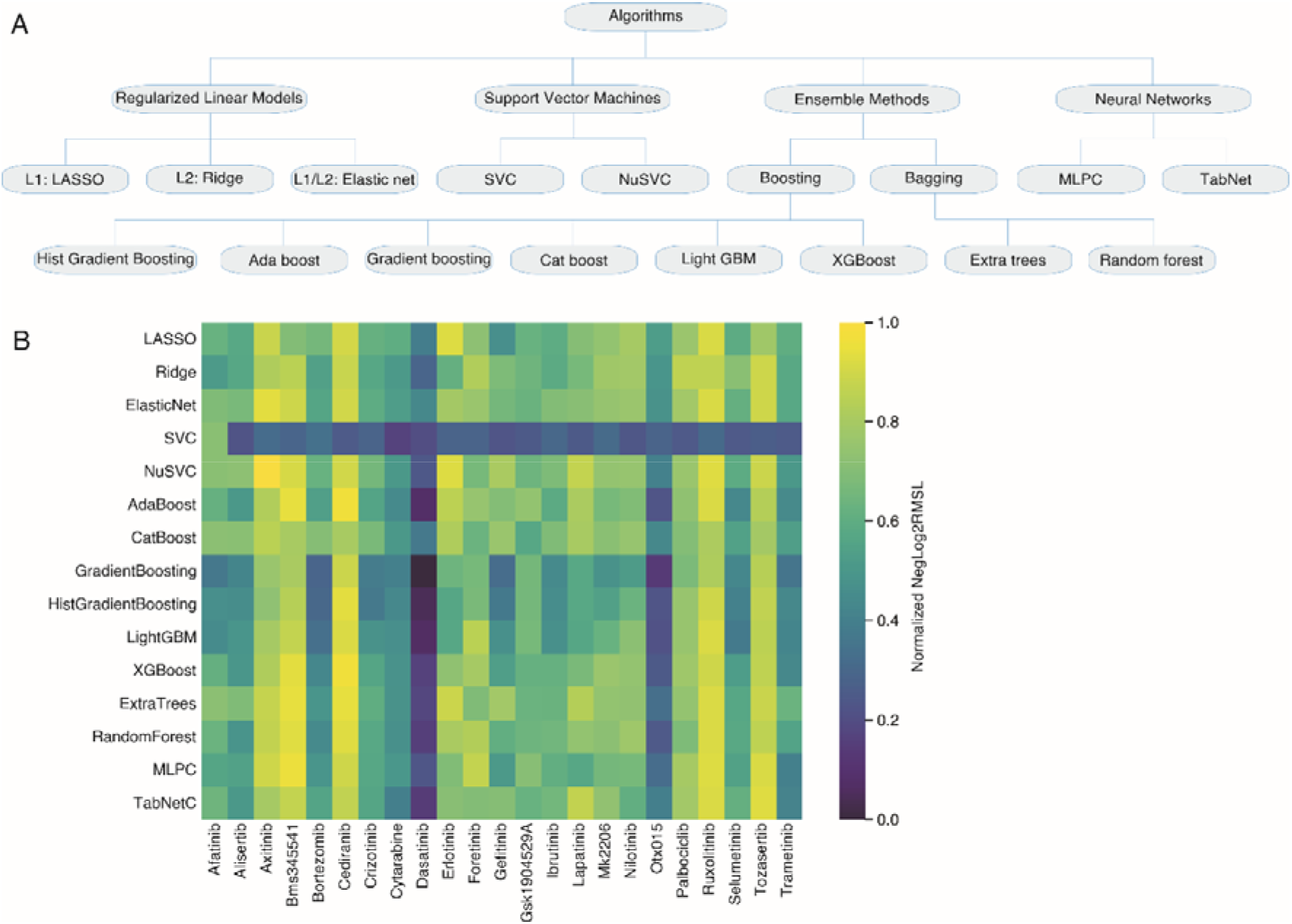
Comparison of different algorithms. (A) A general classification of algorithms used in alphaML platform. (B) AML dataset was used to select features using HVF function. Hyperparameters were optimized using Optuna. The NegLog2RMSL scoring was used to select the best parameters.

Support Vector Machines (SVMs) represent a subset of supervised machine learning methodologies utilized for both classification and regression evaluations of datasets. SVMs operate under the theory of structural risk minimization, aiming to attenuate an upper limit on the potential generalization error as opposed to purely minimizing the error identified during training (Vapnik, 1995). The Support Vector Classification (SVC) is a type of SVM used for classification tasks. In its simplest linear form, the algorithm constructs a hyperplane in a high dimensional feature space to separate the different classes with a maximal margin (Coretes and Vapnik, 1995). This separating hyperplane is influenced by a subset of the initial training data, referred to as support vectors, which are essentially data points in close proximity to the decision boundary. For non-linear applications, SVC integrates the kernel trick to implicitly transform the input data into high-dimensional feature spaces, where a linear delineation is sought after. The nu-Support Vector Classification (NuSVC) signifies another SVM variant, where the problem is parameterized with parameters nu, as opposed to the conventional regularization parameter C seen in SVC. The nu parameter (0 < nu <= 1) symbolizes an upper limit on the training errors proportion, and a lower limit on the fraction of support vectors (Scholkopf, et al., 2000). The NuSVC model provides a way of controlling the number of support vectors and margin errors, thus offering a nuanced control over the model complexity. Both SVC and NuSVC are effective in high-dimensional spaces, even when the number of dimensions exceeds the number of samples, and are versatile in their choice of decision function, which can be linear, polynomial, or a radial basis function.

Ensemble methods combine the predictions of multiple models to create a final prediction that’s more accurate and robust than the individual models alone. The theoretical foundation of these methods is predicated on the amalgamation of weak learners to create a robust learner. The classification of ensemble methods can largely be bifurcated into two principal types: boosting and bagging. Boosting is an ensemble strategy that is designed to evolve a robust classifier from a series of weak classifiers. This is accomplished by building a model predicated on the training data, followed by the creation of a subsequent model which aims to rectify the errors manifested by the former model. Model addition continues either until the training set is flawlessly predicted or until a predefined threshold of models is reached. Adaptive Boosting (AdaBoost) is one of the pioneering boosting algorithms that operates by assigning weights to the observations, attributing higher weightage to challenging instances to classify, and lower to those instances that are classified efficiently (Freund and Schapire, 1997). Gradient Boosting was later proposed, which endeavors to correct the residual errors of the previous predictor by employing gradient descent to minimize the loss when new models are incorporated (Friedman, 2001). Histogram-based Gradient Boosting (HistGradientBoosting) is a variant of gradient boosting which discretizes continuous input features into integer-valued bins, leading to expedited computation and reduced memory footprint (Friedman, 2012). Extreme Gradient Boosting (XGBoost) is a robust and scalable implementation of gradient boosting algorithms, esteemed for its execution velocity and model performance (Chen and Guestrin, 2016). Microsoft Corporation developed the LightGBM, a gradient boosting framework known for its distributed and efficient architecture, superior training speed, efficiency, and capability to manage large-scale data (Ke, et al., 2017). CatBoost, another gradient boosting algorithm, displays exceptional prowess in managing categorical variables (Prokhorenkova, et al., 2018). In contrast to boosting algorithms, bagging (an acronym for bootstrap aggregating) is an ensemble method which involves manipulating the training set through resampling and executing algorithms on it. Bagging diminishes the variance of a base estimator (e.g., a decision tree), by incorporating randomization into its construction procedure and subsequently creating an ensemble from it. The bagging algorithm, Random Forest, merges the concept of bagging and the method of random subspace to fabricate a series of decision trees with regulated variance (Breiman, 2001). The random forest variant extremely randomized trees (Extra Trees) utilizes a random subset of features to bifurcate the data at each node during the construction of the trees (Geurts, et al., 2006).

Artificial Neural Networks (ANNs) are intricate structures comprised of interconnected layers of computational units, often referred to as “neurons”. These units are responsible for the conversion of input data into corresponding outputs. Inter-neuron communication is enabled through connections that facilitate the transfer of signals from one neuron to another. Upon receipt, the signal undergoes processing by the recipient neuron, which subsequently transmits signals to successive neurons in the connection chain. This architecture is hierarchically layered, commencing with an input layer where raw data is introduced. The data then traverses one or more intermediary or “hidden” layers, each responsible for performing specific data transformations, before culminating at the output layer. The multi-layer perceptron classifier (MLPC) is a subset of the broader category of feedforward ANNs (Rumelhart, et al., 1986). MLPs are characterized by a structure comprising a minimum of three node layers: an input layer, a hidden layer, and an output layer. With the exception of the input nodes, each node functions as a neuron employing a nonlinear activation function. For training, MLPs deploy a supervised learning methodology known as backpropagation (Rumelhart, et al., 1986). Lastly, the TabNet algorithm introduces an innovative, high-performance, and interpretable neural network model tailored for tabular data. This model’s distinctive feature is its sequential attention mechanism, which enables the selection of pertinent features at each decision juncture, enhancing its interpretability compared to conventional feed-forward networks or tree-based models. This sequential attention mechanism operates as a learnable mask deployed at each decision point, thus empowering the model to choose the relevant features to examine for each decision (Arik and Pfister, 2021).

We conducted a comparative analysis of all algorithms using 23 inhibitors. The AML dataset was leveraged as a source of feature selection in this particular context. For hyperparameter optimization, we employed the Optuna framework and the ‘custom_score’ to minimize the error. The comparative effectiveness of diverse algorithms and inhibitors was evaluated using the standard measure of normalized NegLog2RMSL. A subset of pharmacological compounds, namely axitinib, BMS-345541, cediranib, ruxolitinib, and tozasertib, demonstrated superior efficacy in most algorithmic interpretations (Figure 5B). While it was observed that ensemble methodologies consistently outperformed others in terms of normalized NegLog2RMSL, algorithms employing regularization techniques also yielded commendable outcomes, underlining the relevance of regularization in processing high-dimensional datasets.

### The interpretive module of the model

The preponderance of algorithmic configurations within the alphaML framework encompasses innate methodologies for deducing feature importance. However, certain algorithms, namely linear algorithms, SVCs and MLPC are devoid of such integral functionality. Consequently, in addition to these inherent methodologies, we have assimilated permutation feature importance from sklearn and global feature importance from Shapley Additive Explanations (SHAP), which possess the capability of discerning influential features, independent of their algorithmic origins (Breiman, 2001; Lundberg, et al., 2020; Strobl, et al., 2008). SHAP is a cohesive measure of feature significance, attributing an importance value to each feature for a specific prediction. This measure finds its roots in the Shapley values concept, a derivative of cooperative game theory that assures equitable distribution of payoffs among players depending on their contribution to the total payoff (Lundberg and Lee, 2017). SHAP values provide an interpretation of the influence of a specific value for a particular feature compared to the prediction we would procure if that feature assumed a baseline value. Essentially, SHAP attributes an importance value to each feature for a specific prediction, symbolizing the degree to which the feature’s value contributes to the deviation of the prediction from the baseline prediction. SHAP proposes a methodology to calculate universal feature significance by averaging the absolute SHAP values for each feature across all instances within a dataset. This provides an understanding of the effect of each feature on model prediction across numerous instances, as opposed to a singular instance. Features are then typically stratified by these universal importance values, providing a perspective on which features are predominantly influential in the model’s predictions on average. Hence, SHAP can be employed to discern both local and universal feature significance.

Additionally, we have assimilated the Local Interpretable Model-agnostic Explanations (LIME), a technique devised for elucidating the predictions of any machine learning classifier (Ribeiro, et al., 2016). LIME approximates the decision boundary of the complex model locally with a simplified model, such as a linear model, which is easier to interpret. For a specific instance, LIME generates a new dataset of perturbed samples, procures the predictions of these samples using the complex model, and then fits the simplified model on this new dataset. The weights of the simplified model are learned based on the proximity of each perturbed sample to the instance of interest. The coefficients of the simplified model serve as an explanation of the contribution of each feature to the prediction for the instance of interest. The advantage of LIME is that it offers local interpretability, elucidating the model’s decisions on a per-instance basis. However, it can also be employed to evaluate universal feature significance by aggregating local explanations for a representative set of instances.

The evaluation of model efficacy was conducted through the application of test data and the implementation of a 5-fold repeated cross-validation. Figure 6 illustrates data derived specifically from the application of crizotinib and the XGBoost algorithm. The construction of the confusion matrix (Figure 6A) was achieved using test data, while the receiver operating characteristic area under the curve (ROC-AUC) (Figure 6B) and accuracy plot (Figure 6C) were generated through the use of 5-fold repeated cross-validation. Additionally, the computation of scores for different metrics (Figure 6D) was conducted for the test sample. The test samples were further utilized for the ascertainment of feature importance from the model (Figure 6E), the permutation (Figure 6F), and the SHAP (Figure 6G). Feature importance for individual test samples was depicted utilizing both SHAP and LIME, with exemplification through a crizotinib sensitive sample (Figure 6H and 6J) and a crizotinib resistant sample (Figure 6I and 6K). Interestingly, a recurring pattern was observed wherein the expression of SIGLEC1 and CHIT1 appeared to exert influence on the prediction outcomes.

**Figure 6.**
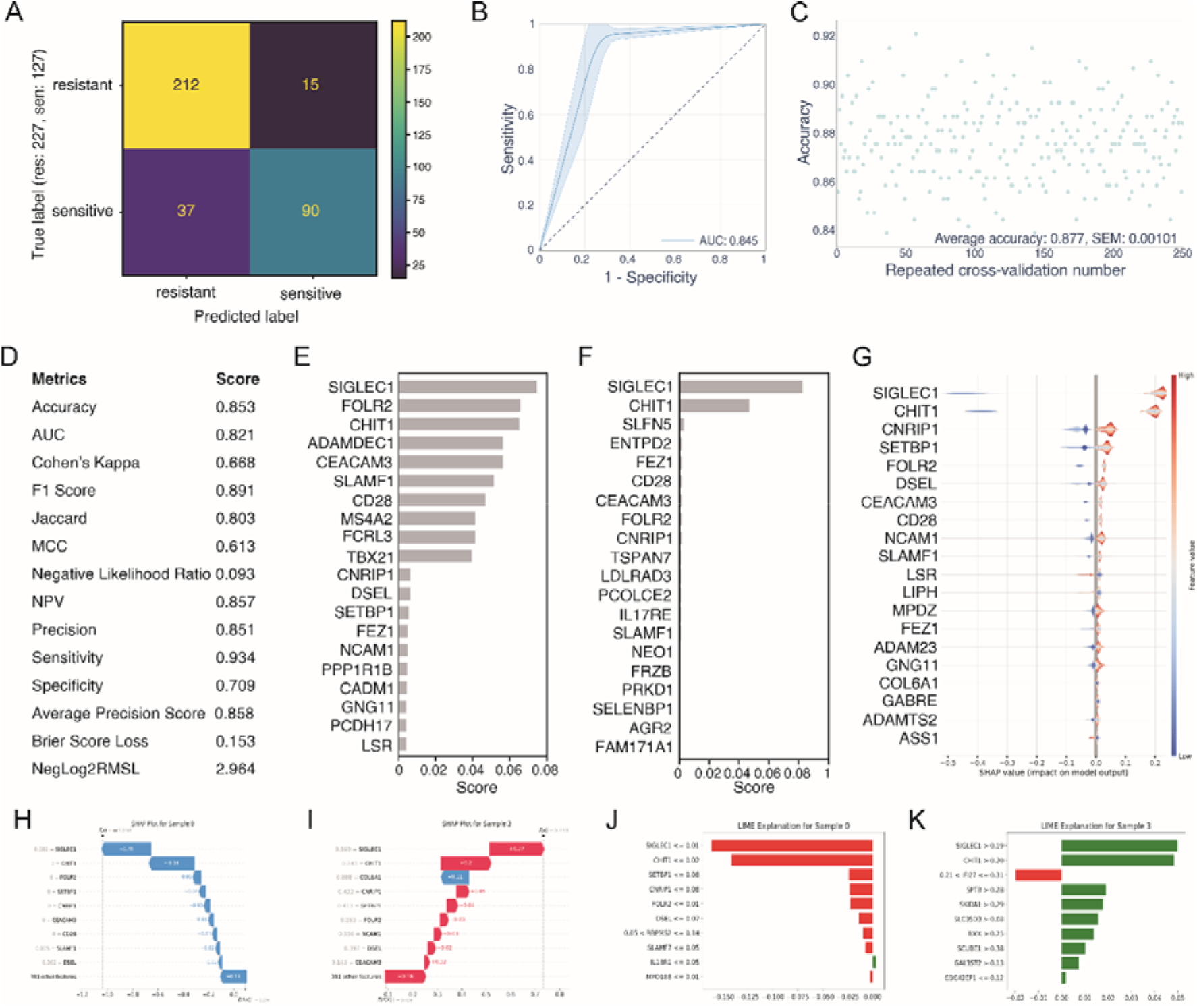
Evaluation and Interpretation of Predictive Model. The dataset comprised of predictive and explanatory data derived from crizotinib and the XGBoost algorithm. (A) A confusion matrix was constructed utilizing the test sample dataset. (B) The ROC-AUC was generated through a 5-fold CV approach. (C) Average precision was ascertained via a 5-fold CV technique. (D) Various metric scores were computed utilizing the test sample data. (E) Feature significance was established using the inherent function of XGBoost, wherein the top 20 prominent features were illustrated. (F) A Permutation Feature Importance method was employed to identify key features. (G) Global SHAP values were charted for the test samples. (H) A SHAP local explanation was provided for the sample predicted as sensitive. (I) A SHAP local explanation was presented for the sample predicted as resistant. (J) An explanation using LIME was provided for the sample anticipated as sensitive. (K) A LIME explanation was delineated for the sample expected to be resistant.

### Prediction module

The prediction module integrated within the computational framework operates as an autonomous unit, maintaining independence from the model construction platform. This prognostic component accepts a trained algorithm and corresponding feature set utilized during the model training phase, enabling predictive operations and elucidation of the prediction rationale via SHAP and LIME, utilizing data as an input source. RNAseq data from 3071 AML patients were procured from the Genomic Data Commons (GDC) data portal (portal.gdc.cancer.gov) for this purpose. The samples demonstrated an amplified sensitivity towards bortezomib, a proteasome inhibitor, foretinib, a multi-targeted tyrosine kinase inhibitor, and trametinib, a MEK inhibitor, (Figure 7A) thereby underscoring the potential of a predictive model in identifying patient-specific therapeutic strategies. In addition, the global feature significance as indicated by SHAP (Figure 7B) and local importance as defined by LIME (Figure 7C and 7D) provided valuable insights into influential regulators, with crizotinib serving as a prime example in this context.

**Figure 7:**
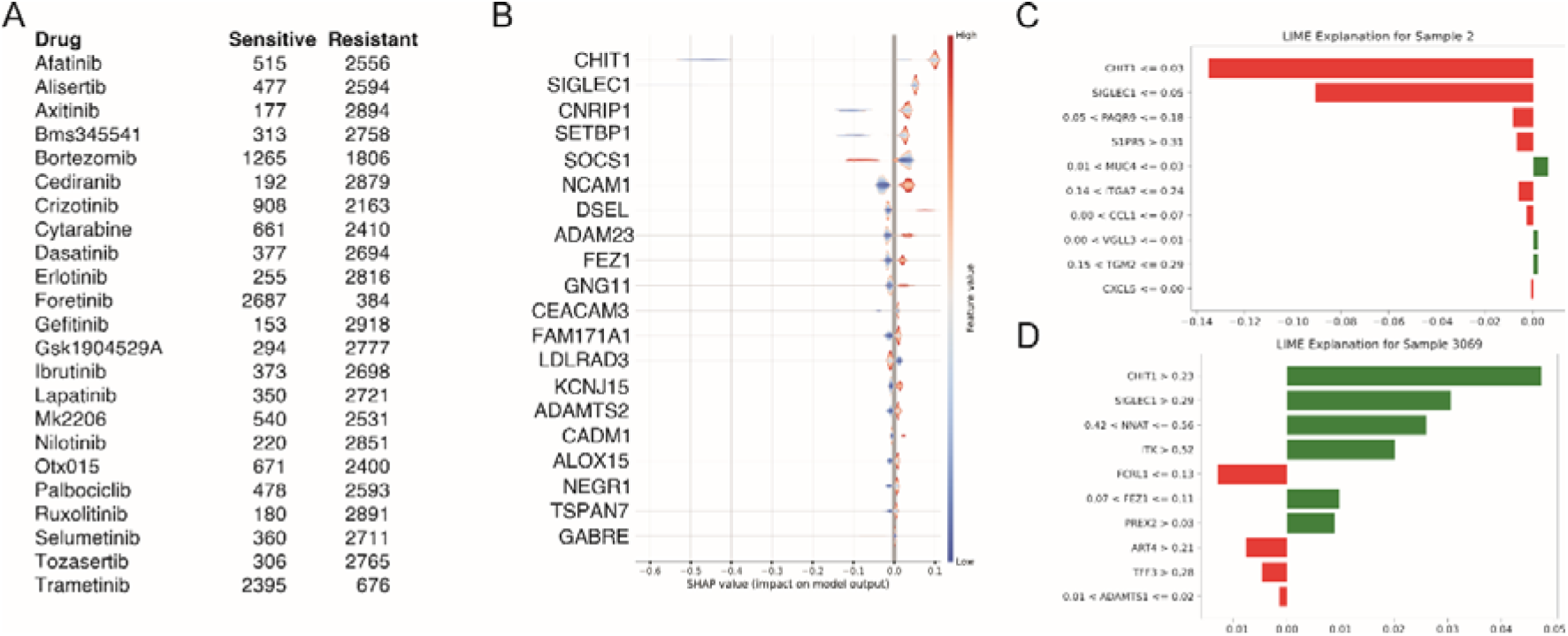
Inferential analysis with XGBoost algorithms. AML patient dataset was employed to forecast the responsiveness towards 23 therapeutic drugs. (A) Enumeration of samples deemed sensitive or resistant as discerned by the XGBoost algorithm. (B) Comprehensive SHAP illustration associated with the crizotinib prediction. (C) LIME elucidation corresponding to the sample classified as sensitive. (D) LIME interpretation pertinent to the sample designated as resistant.

### Versatility of the platform

To evaluate the robustness and applicability of our platform across diverse domains, we deployed it on six distinct datasets, namely: Bank Marketing, Credit Card Fraud Detection, Pima Indians Diabetes, NATICUSdroid (Android Permissions), Student Dropout and Academic Success prediction, and Breast Cancer Wisconsin (Diagnostic) dataset. The first five datasets were utilized to ascertain the consistency of the model’s predictive performance between the training and testing subsets, as implemented through the XGBoost algorithm (Figure 8A). For the final dataset - Breast Cancer Wisconsin (Diagnostic) - we utilized it to generate a confusion matrix (Figure 8B) and ROC-AUC curve (Figure 8C), repeated accuracy plot (Figure 8D). The cumulative evidence affirms the broad-spectrum utility of our proposed platform.

**Figure 8:**
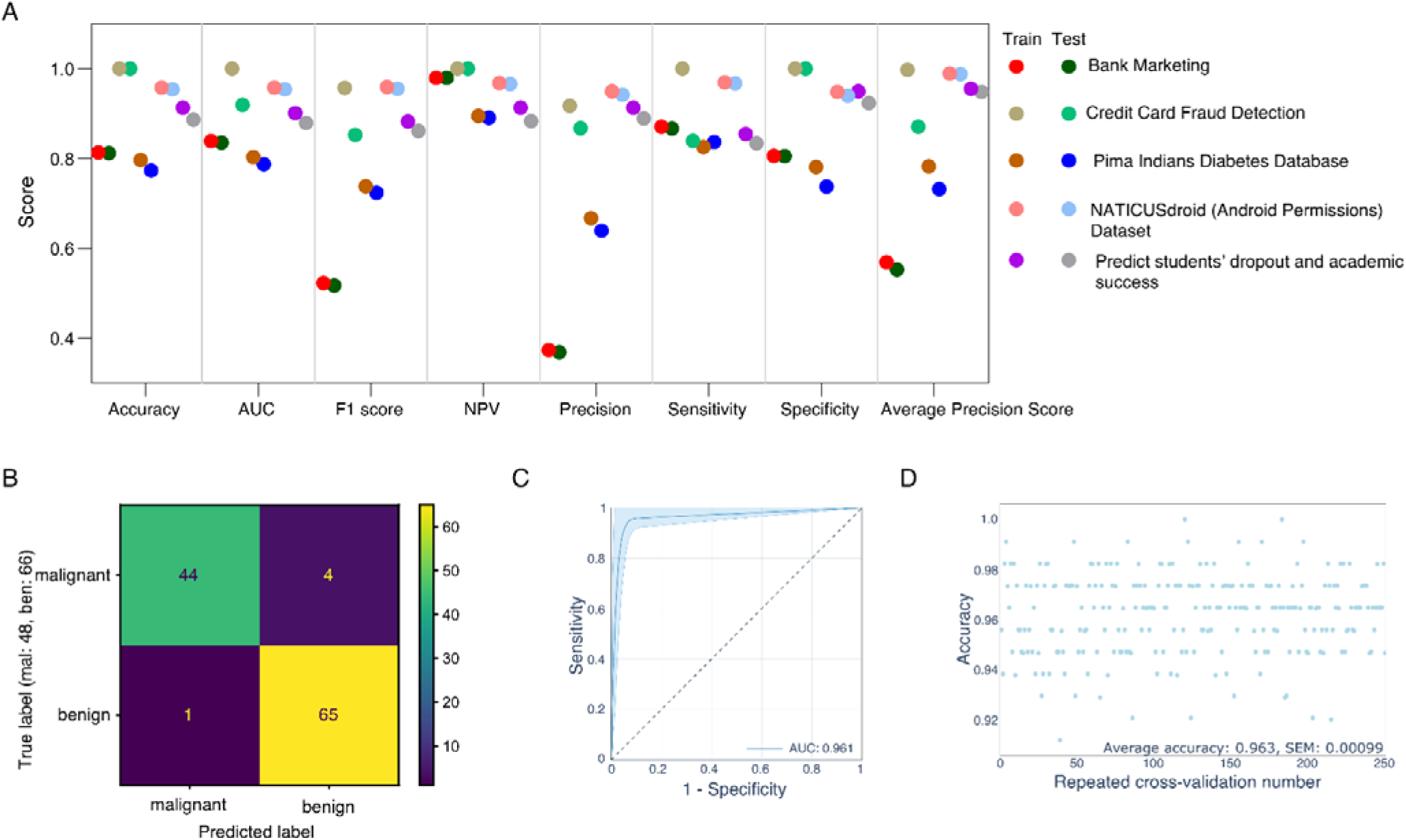
Evaluation of model adaptability across diverse datasets. To assess the platform’s versatility, we tested it against six disparate datasets. (A) Performance metrics were calculated utilizing the XGBoost algorithm, with hyperparameter tuning executed through Optuna. All attributes encompassed in these datasets were considered. (B-D) Utilizing the Wisconsin Breast Cancer (Diagnostic) dataset, the model was constructed and validated employing the XGBoost framework, with Optuna facilitating hyperparameter optimization.

## Discussion

In the presented research, we delineated a facile binary classification framework aimed at predicting therapeutic response in oncology, employing advanced machine learning techniques. This versatile platform integrates an assortment of critical components, such as feature selection, data normalization, data sampling, hyperparameter optimization, model training and validation, testing, and model interpretability. The assemblage of these modules enables a comprehensive machine learning solution. Its potency was validated by employing pharmacogenomic data across a broad spectrum of drugs and diseases, signifying its wide applicability to various tabular data formats.

The feature selection module is operable as an autonomous entity, and its efficacy was evaluated across a variety of disease classes utilizing 23 distinct pharmacological compounds. The results emphatically illustrate that the selected features manifest disease-specific as well as drug-specific traits, thereby inferring the indispensability of predictive models tailored to specific pathologies and pharmaceutical agents. Additionally, through the overexpression of BEX1 in ovarian cancer cells, partial validation of the utility of the feature selection component has been achieved.

The imbalanced-learn library was employed within the alphaML framework to govern undersampling and oversampling applications. Interestingly, the results indicated a preference for the ‘no sampling’ approach over traditional methods in analyzing pharmacogenomic data. The observations suggest that preserving the inherent major and minor class proportions in the dataset yielded superior performance, even as the class imbalance parameter was triggered during the model’s construction phase. However, the research also recognizes that the comparative performance of these sampling methods could vary significantly based on the problem at hand, highlighting the constant requirement for comprehensive comparative assessments. Test data integrity was ensured via pre-sampling partitioning, thereby eliminating any potential sampling bias.

The Optuna hyperparameter optimization technique was employed to assess its efficacy, using three performance indices: Hamming loss, kappa_mcc_error, and a newly introduced metric dubbed ‘custom_score’. The Hamming loss and kappa_mcc_error indices use validation data to steer the model’s optimization, whereas the custom_score metric utilizes both training and validation datasets to bridge performance gaps. When integrated with the Optuna scoring framework, the custom_score metric appeared to augment performance, as indicated by the superior NegLog2RMSL metrics in the study of 23 inhibitors across four disease models. A comparative analysis of different algorithms and search methods revealed that combining custom_score with Optuna yielded the most favorable NegLog2RMSL, signifying that the incorporation of the custom_score metric may potentially enhance binary classification model performance. This research featured an exhaustive comparison of various algorithms using 23 inhibitors, sourcing feature selection from the AML dataset. The Optuna framework was employed for hyperparameter optimization, alongside a ‘custom_score’ to curtail errors.

Normalized NegLog2RMSL was used as the benchmark for comparative performance assessment, revealing superior efficiency of certain drugs such as axitinib, BMS-345541, cediranib, ruxolitinib, and tozasertib. The results underscored the commendable performance of ensemble methods and the significance of regularization techniques when dealing with high-dimensional datasets. Model evaluation was performed using test data and 5-fold repeated cross-validation, as depicted in the data derived from the application of crizotinib and the XGBoost algorithm. Metrics such as the confusion matrix, ROC-AUC, accuracy plot, and various other metric scores were calculated and visualized, providing diverse insights into the model’s performance and efficacy. Additionally, test samples were used to elucidate the model’s feature importance, as illustrated by permutations and SHAP. The recurrent pattern of SIGLEC1 and CHIT1 expression impacting prediction outcomes was notable, thereby indicating potential predictive value of these biomarkers.

The prediction module, a standalone component of the computational framework, functions independently of the model construction platform. It ingests a pre-trained algorithm and corresponding feature set, thus enabling predictive functionality and the explanation of prediction rationales via SHAP and LIME, utilizing RNAseq data from 3071 AML patients procured from the GDC data portal. The sensitivity displayed by the patient samples towards bortezomib, foretinib, and trametinib underscores the potential of predictive models in devising patient-specific therapeutic strategies. Moreover, the global and local feature importance as delineated by SHAP and LIME, respectively, offer insights into influential regulators, exemplified by crizotinib.

In conclusion, this study effectively demonstrates the potential of a comprehensive machine learning framework in optimizing therapeutic strategies for oncology patients. Through the integrated analysis of complex pharmacogenomic data, this approach not only highlights the value of tailored disease and drug-specific predictive models, but also underscores the promising role of computational methodologies in advancing personalized medicine.

## Availability

The Python scripts are accessible to individuals possessing fundamental computational competencies and rudimentary proficiency in the management of tabular data via applications akin to Microsoft Excel. For enhanced user interaction, we furnish a Graphical User Interface (GUI). The Python library, alphaML, can be procured from the Python Package Index (PyPI) at www.pypi.org/project/alphaml/, or directly from the GitHub repository at www.github.com/kazilab/alphaML.

## Data Procurement

The source of annotated data was derived from the scDEAL and BeatAML research studies (Bottomly, et al., 2022; Chen, et al., 2022). The dataset utilized for attribute selection was procured from The Cancer Genome Atlas (TCGA) and the Therapeutically Applicable Research to Generate Effective Treatments (TARGET) databases. The AML data employed for prognostic analyses was obtained from the GDC data portal (portal.gdc.cancer.gov).

## Supplementary Materials

Procedure for configuring the alphaML platform within a Python-based computational environment:

### Installation

1. *Install Anaconda*: Download the latest version of Anaconda from www.anaconda.com. Install by double clinking the downloaded installer. When the installer asks for the installation type, choose “Just Me”.
2. *Install alphaml*: For Windows - search for *Anaconda Powershell Prompt in Windows search and click on it*. Type the following command and press “Enter” to install alphaml and dependent packages. `pip install alphaml` For Mac - search for Terminal and click on it. Before installing the package, activate the base conda environment by typing the following command and pressing “Enter”: `conda activate` “base” should be prepended, then type the following command and press “Enter” to install alphaml: `pip install alphaml` After installation, deactivate the conda environment by the following command: `conda deactivate` Alternatively, Anaconda Navigator can be used (platform independent). Launch Anaconda Navigator and from the navigator launch JupyterLab. It will open a browser window. Click on Terminal from the browser window. Type the following command in Terminal/PowerShell window and press “Enter” to install alphaml: `pip install alphaml`

### Run alphaml

1. In Windows, search for “Anaconda Powershell Prompt” in the Windows search bar and click on it. Type the following command and press “Enter” to open the alphaML GUI: `python -c “from alphaml import guir”`
2. In Mac, open Terminal. First, activate the base conda environment with the following command: `conda activate` After seeing “base” prepended to the prompt, run the alphaML GUI with the following command: `python -c “from alphaml import guir”` After use, deactivate the conda environment by typing the following command: `conda deactivate`
3. If Anaconda Navigator is being used, launch it and then launch JupyterLab from within Navigator. This will open a browser window. Click on “Terminal” in the browser window, then type the following command and press “Enter” to open the alphaML GUI: `python -c “from alphaml import guir”`

### Using alphaML from the GUI

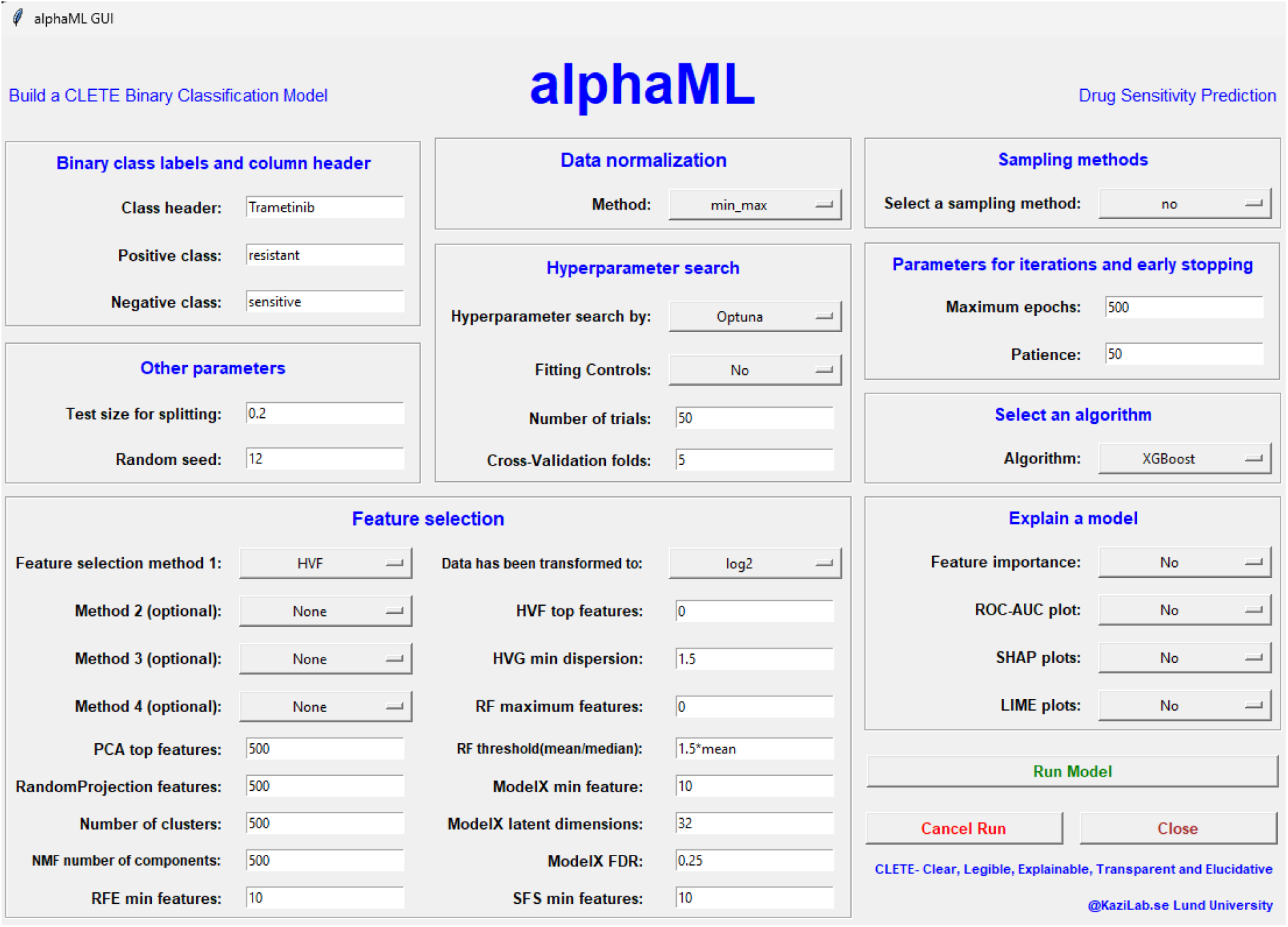

1. Launch the alphaML GUI. Press the “Run Model” button. This will automatically generate two folders, namely “alphaML_data” and “alphaML_results”, in the current user’s Documents directory.
2. The “alphaML_data” folder should contain at least five files named “data_for_feature_selection.csv”, “data_labels.csv”, “labeled_data.csv”, “suggested_features.csv”, and “unlabeled_data.csv”. alphaML first checks the presence of these files and, if they are missing, it will attempt to download sample data. Please ensure that these filenames are unaltered, as they have been hardcoded into alphaML.
3. The “labeled_data.csv”, “data_for_feature_selection.csv”, and “unlabeled_data.csv” files should all contain tabular data. The first column of these tables should include sample names, and each row must represent a distinct sample. Ensure the column headers denote feature names. The “data_labels.csv” file should contain binary labels corresponding to the labeled data. The “suggested_features.csv” file, a single-column file, is designed to accept pre-selected features for model input. Please note: The “labeled_data.csv” file is considered as the annotated data which has corresponding binary labels in the “data_labels.csv”. Example: “labeled_data.csv”.

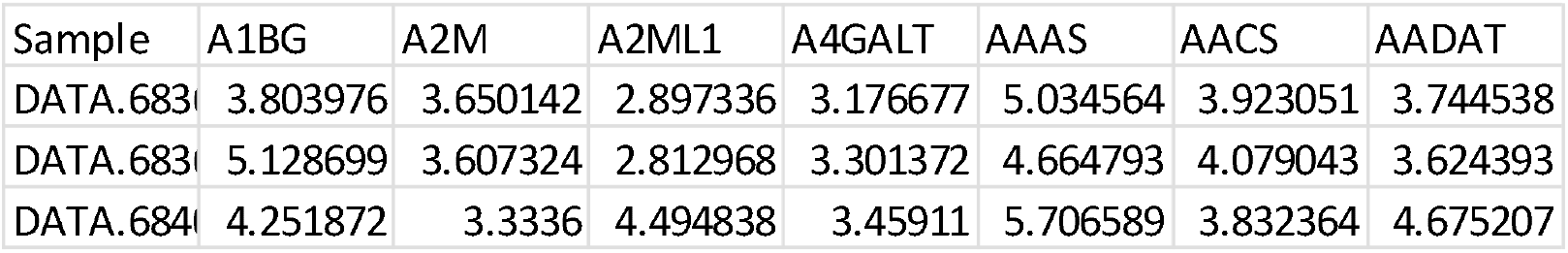 Example: “data_labels.csv “

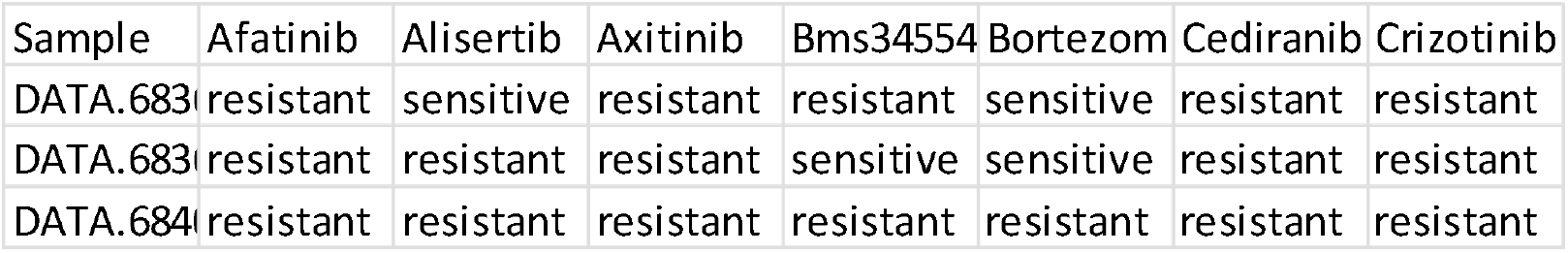 The “data_for_feature_selection.csv” file should contain the data you intend to use for feature selection. This can include specific data like that related to a particular disease. If you don’t have specific data for this purpose, you can use your labeled data. In this case, copy the contents of your “labeled_data.csv” file into the “data_for_feature_selection.csv” file. Example: “data_for_feature_selection.csv”

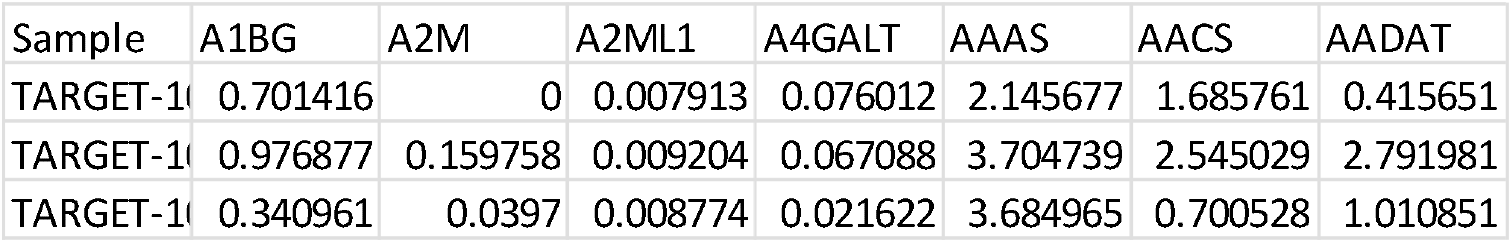 The “unlabeled_data.csv” file should contain the unannotated data, which lacks binary labels. If you don’t have a separate set of unannotated data, you can use your labeled data. In this case, copy the contents of your “labeled_data.csv” file into the “unlabeled_data.csv” file. Example: “unlabeled_data.csv”

4. *Binary class labels and column header:* The “Class header” is used to find the input string in data_label.csv for information about the binary classes. Input in the “Class header” must be identical to the respective class header in “data_labels.csv”. The “data_labels.csv” requires at least two columns: “Sample” and a column with class information. The “Positive class” represents the outcome or event of interest for prediction, such as resistance, cancer, diabetes, etc. Conversely, the “Negative class” represents the other possible outcome or event, such as sensitivity, healthy control, etc. The binary class label converter in alphaML removes empty spaces and converts labels to lowercase before encoding the “Positive class” label as “1” and the “Negative Class” label as “0”. For instance, “downregulated” or “Downregulated” or “Down regulated” or “Down Regulated” will all be encoded as the same class.

|  | A1BG | A2M | A2ML1 | A4GALT | AAAS | AACS | AADAT |
| --- | --- | --- | --- | --- | --- | --- | --- |
| aml_ohsu | 0.330713 | 0.280005 | 0.02153 | 0.520474 | 5.475134 | 2.719014 | 0.284122 |
| aml_ohsu | 0.333274 | 1.813633 | 0.05036 | 4.993026 | 5.252081 | 2.81072 | 0.212554 |
| aml_ohsu | 0.310227 | 0.609844 | 0.087837 | 3.126578 | 5.579161 | 2.789831 | 0.192626 |
| aml_ohsu | 0.029552 | 2.409636 | 0.08932 | 1.253376 | 5.486577 | 3.227344 | 1.1814 |

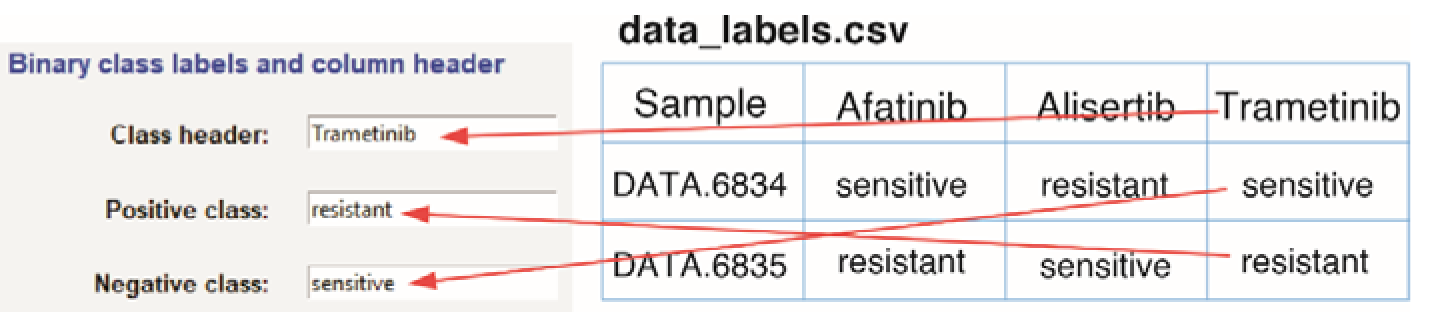
5. *Other parameters:* The parameter “Test size for splitting” determines the division of samples for model building and testing. By default, the ratio of training to test samples is 80:20, represented by 0.2 (test sample size). Modify this parameter between 0.05 and 0.5 to alter the test sample size. The “Random seed” should be a positive number.
6. *Feature selection module:* This module extracts important features or loads predefined features for model development. The feature selection module in alphaML provides access to a panel of feature selection methods. Users can use a single feature selection method from the drop-down list or use up to four methods in combination. Method 1 is mandator and from method 2, 3 and 4, user can select any or none of them. *PCA top features:* Used by SelByPCA (select by PCA) method and can be between 1 and the initial number of features. *Random Projection features:* Used by RandomProjection method and can be between 1 and the initial number of features. *Number of clusters:* Used by SelByClustering (select by clustering) method and can be between 1 and the initial number of features. *NMF number of components:* Used by SelByNMF (select by NMF) method and can be between 1 and the initial number of features. *RFE number of components:* Used by RecursiveFeatElim (Recursive feature elimination) method and can be between 1 and the initial number of features. Computationally expensive method and is not recommended for a large number of initial features. *Data has been transformed to:* Used by HVF (highly variable features) method. HVF converts log transformed data to anti-log, therefore, it is important to mention here what type of log transformation has been used in data appended in “data_for_feature_selection.csv” file. Select from a drop-down list. Select None if data has not been transformed to log. *HVF top features:* Used by HVF (highly variable features). The number of features to be selected. Input can be between 1 and the initial number of features. If “0”, this option will be ignored. *HVF Min dispersion:* Used by HVF (highly variable features). The number of features that pass normalized dispersion value. Input can be any positive number. Only used if *HVF top features* is “0”, otherwise, this option will be ignored. *RF Maximum features:* Used by SelectByRF (select by random forest). The maximum number of features to be selected that pass “RF threshold”. If “0”, this option will be ignored. *RF Threshold:* Used by SelectByRF (select by random forest). To select features that pass a threshold. It can be mean or median and multiplied by a positive number. *ModelX Min features:* Used by ModelX (Model-X knockoffs). The suggested minimum number that should be passed to proceed with the selected features. If ModelX identifies less features than the “*ModelX Min features”* the alphaML feature selection module will ignore this method and continue with the input features. *ModelX Latent dimension:* Used by ModelX (Model-X knockoffs). Latent dimensions to be used by autoencoder incorporated in ModelX method. *ModelX FDR:* Used by ModelX (Model-X knockoffs). False discover rate cut-off to be used by ModelX. *SFS Min features:* Used by SeqFeatSel (Sequential Feature selection) method and can be between 1 and the initial number of features. Computationally expensive method and is not recommended for a large number of initial features. *“RemoveHighCorrFeat” method in Method 3 and Method 4 can be applied to remove feature that correlates more than 90% with another feature*. *“IterativeFeatSel” method in Method 3 and Method 4 can be applied to select 1 to 3 most important features. Computationally extremely expensive for a large number of initial features*. *“None” method in Method 2, Method 3 and Method 4 can be used to ignore a method*. *Feature selection module can be used independently by selecting. “*
7. Data Normalization: Two data normalization options are available: “min_max” and “standardization”. If no normalization is required, select “None” from the dropdown list.
8. *Hyperparameter search:* Optuna, Bayes, or Grid search methods can be selected from the dropdown menu. These represent Optuna, BayesSearchCV, and GridSearchCV respectively. A “Predefined” option, where all parameters are hardcoded, is also available. *Fitting Controls:* This option is applicable only for “Optuna”. From a drop-down menu if *“Yes”* is selected, Optuna will use a custom scoring method that considers train metrics to control overfitting. *Number of trials:* How many trials are to be performed for hyperparameter search. *Cross-validation folds:* Number of cross validations to divide data.
9. *Sampling methods:* By default, alphaML does not apply any sampling method. However, “under” and “over” sampling methods can be selected from the dropdown menu.
10. *Parameters for iterations and early stopping:* Used by TabNet only.
11. *Select an algorithm:* An algorithm can be chosen from 15 different options available in a dropdown menu. Alternatively, a quick test can be performed by selecting “Test_Briefly”. If “None” is selected, no model will be built.
12. Explain a model: To perform any or all of four methods, including “Feature importance”, “ROC-AUC plot”, “SHAP plots”, and “LIME plots”, select from dropdown menus.
13. To initiate the run, click on the “Run Model” button. Click on the “Cancel Run” button to abort a run and the “Close” button to close the GUI window.

### The alphaML result files

1. Depending on how many feature selection methods have been selected, result folder can contain 1-4 CSV files with name as follows: ClassHeader_Method1_None_ None _ None_Feat_M1.csv ClassHeader_Method1_Method2_ None_ None_Feat_M2.csv ClassHeader_Method1_Method2_Method3_ None_Feat_M3.csv ClassHeader_Method1_Method2_Method3_Method4_Feat_M4.csv
2. Two CSV files containing normalized labeled and unlabeled data. ClassHeader_Method1_normalized_labeled_data.csv ClassHeader_Method1_normalized_unlabeled_data.csv
3. A PDF file containing parameters identified by hyperparameter search. ClassHeader_SamplingMethod_HyperparameterSearch_Algorithm_fit_FittingControl_parameters.pdf
4. The model file that can be used for prediction. ClassHeader_SamplingMethod_HyperparameterSearch_Algorithm_fit_FittingControl_model.pkl
5. An accuracy and loss plot showing both training and test scores. ClassHeader_SamplingMethod_HyperparameterSearch_Algorithm_fit_FittingControl_valid_accuracy_curve.pdf
6. A PDF file containing the confusion matrix created using test samples. ClassHeader_SamplingMethod_HyperparameterSearch_Algorithm_fit_FittingControl_confusion_matrix.pdf
7. A CSV file containing train and test scores. ClassHeader_SamplingMethod_HyperparameterSearch_Algorithm_fit_FittingControl_scores.csv
8. An XLSX file containing true test labels, prediction labels and prediction probability. ClassHeader_SamplingMethod_HyperparameterSearch_Algorithm_fit_FittingControl_ test_prediction.xlsx
9. A CSV file containing feature importance calculated by the model (if available) and permutation importance. ClassHeader_SamplingMethod_HyperparameterSearch_Algorithm_fit_FittingControl_global_feat_imp.csv
10. A PDF file containing global SHAP plots and SHAP plots for each test sample. ClassHeader_SamplingMethod_HyperparameterSearch_Algorithm_fit_FittingControl_ test_SHAP.pdf
11. A PDF file containing LIME plots for each test sample. ClassHeader_SamplingMethod_HyperparameterSearch_Algorithm_fit_FittingControl_ test_LIME.pdf
12. A CSV file containing avu, fpr, tpr, accuracy and thresholds for repeated k-fold analysis. ClassHeader_SamplingMethod_HyperparameterSearch_Algorithm_fit_FittingControl_repeated_accuracy.csv
13. A PDF file containing ROC-AUC curve from repeated k-fold analysis. ClassHeader_SamplingMethod_HyperparameterSearch_Algorithm_fit_FittingControl_ AUC_curve.pdf
14. A PDF file containing dot plot of accuracy measurements from repeated k-fold analysis. ClassHeader_SamplingMethod_HyperparameterSearch_Algorithm_fit_FittingControl_accuracy_plot_k_fold_cv_roc.pdf
15. A text file containing log. alphaML_run.log
16. If Optuna is used for hyperparameter search, and additional PDF file with hyperparameter importance will be created. ClassHeader_SamplingMethod_HyperparameterSearch_Algorithm_fit_FittingControl_optuna_param_importance.pdf

### Run a prediction using alphaPred

1. In Windows, search for *Anaconda Powershell Prompt in Windows search and click on it*. Type the following command and press “Enter” to open alphaPred GUI. `python -c “from alphaml import guip”`
2. In Mac, search for Terminal and click on it and type the following command and press “Enter”. `conda activate` “base” should be prepended, then type the following command and press “Enter” to open alphaPred GUI. `python -c “from alphaml import guip”` After use deactivate the conda environment by the following command. `conda deactivate`
3. If Anaconda Navigator is being used, Launch Anaconda Navigator and from the navigator Launch JupyterLab. It will open a browser window. Click on Terminal from the browser window. Type the following command in Terminal/PowerShell window and press “Enter” to open alphaPred GUI. `python -c “from alphaml import guip”`

### Using alphaPred from the GUI

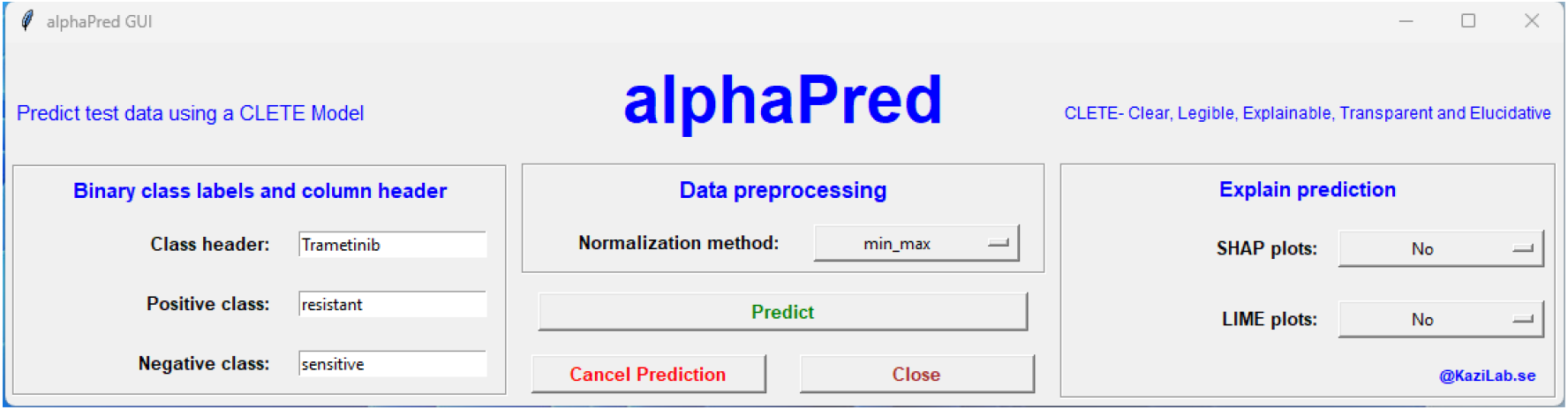

1. Once alphaPred GUI is opened, press on “Predict”. The alphaPred will create two folders (alphaPred_data and alphaPred_results) in current users Documents folder.
2. The alphaML_data folder should contain at least three files with names “model.pkl”, “selected_features.csv”, and “test_data.csv”. The alphaPred first checks the availability of these three files, otherwise, attempts downloading example data. Names of all three files have been hard coded in alphaPred, so must be kept the same.
3. The “model.pkl” is the model file built by alphaML to be used for prediction. The model file should be copied from alphaML_results and renamed.
4. The “selected_features.csv” contains features used to build the model. This file should be copied alphaML_results and renamed. Example: “selected_features.csv”

5. The “test_data.csv” contains tabular data for prediction. The first column should contain sample names and each row should represent a sample. Column headers must represent feature names. Example: “test_data.csv”.

|  |
| --- |
| Highly Var |
| ABCA13 |
| ABCA2 |
| ABTB2 |
| ACOT11 |

6. *Binary class labels and column header:* The “Class header” does not have any role in prediction but can be useful for documentation. The “Positive class” and “Negative class” names are used to convert “1” and “0” to class labels.
7. *Data preprocessing:* The same preprocessing method must be applied that used for alphaML model building.
8. *Explain prediction:* SHAP plots and LIME plots can be generated by selecting “Yes” from the drop-down menu.
9. Clicking on the “Predict” button will initiate the prediction. The “Cancel Run” button will abort a run and click on “Close” button to close the GUI window.

| Sample | A1BG | A2M | A2ML1 | A4GALT | AAAS | AACS | AADAT |
| --- | --- | --- | --- | --- | --- | --- | --- |
| TARGET-1 | 0.701416 | 0 | 0.007913 | 0.076012 | 2.145677 | 1.685761 | 0.415651 |
| TARGET-1 | 0.976877 | 0.159758 | 0.009204 | 0.067088 | 3.704739 | 2.545029 | 2.791981 |
| TARGET-1 | 0.340961 | 0.0397 | 0.008774 | 0.021622 | 3.684965 | 0.700528 | 1.010851 |
| TARGET-1 | 0.327687 | 0.050467 | 0 | 0.036609 | 3.746033 | 1.06757 | 0.122871 |
| TARGET-1 | 0.201885 | 0.3744 | 0.033511 | 0.1595 | 1.239398 | 0.983969 | 2.133892 |

### The alphaPred result files

1. A CSV file containing normalized test data. ClassHeader__Algorithm_fit_normalized_test_data.csv
2. An XLSX file containing sample name, prediction class and probability scores. ClassHeader__Algorithm_fit__test_prediction.xlsx
3. Two PDF files with SHAP and LIME plots. ClassHeader__Algorithm_fit__test_SHAP.pdf ClassHeader__Algorithm_fit__test_LIME.pdf
4. A text file containing log. alphaPred_run.log

